# Evaluating the Impact of Purifying Selection on Species-level Molecular Dating

**DOI:** 10.1101/622209

**Authors:** Chong He, Dan Liang, Peng Zhang

**Affiliations:** State Key Laboratory of Biocontrol, College of Ecology and Evolution, School of Life Sciences, Sun Yat-Sen University, Guangzhou, China

**Keywords:** Purifying selection, molecular clock, neutral theory, molecular dating, rate of evolution

## Abstract

The neutral theory of molecular evolution suggests that the constancy of the molecular clock relies on the neutral condition. Thus, purifying selection, the most common type of natural selection, could influence the constancy of the molecular clock, and the use of genes/sites under purifying selection may produce less reliable molecular dating results. However, in current practices of species-level molecular dating, some researchers prefer to select slowly evolving genes/sites to avoid the potential impact of substitution saturation. These genes/sites are generally under a strong influence of purifying selection. Here, from the data of 23 published mammal genomes, we constructed datasets under various selective constraints. We compared the differences in branch lengths and time estimates among these datasets to investigate the impact of purifying selection on species-level molecular dating. We found that as the selective constraint increases, terminal branches are extended, which introduces biases into the result of species-level molecular dating. This result suggests that in species-level molecular dating, the impact of purifying selection should be taken into consideration, and researchers should be more cautious with the use of genes/sites under purifying selection.

## Introduction

The foundation of molecular dating lies in the molecular clock phenomenon discovered in the 1960s (Margoliash 1963; Zuckerkandl and Pauling 1965). The theoretical population geneticist, Motoo Kimura, noted that the neutral theory can provide an explanation for the molecular clock phenomenon (Ohta and Kimura 1971; Kimura 1977; Takahata 1987; Ohta 1992; Takahata 2007; Nei et al. 2010). This viewpoint about the molecular clock is based on a well-known conclusion of the neutral theory that the substitution rate under selective neutrality is expected to be equal to the mutation rate (Kimura 1983; Ohta 1992; Nei et al. 2010).

First, neutral theory suggests that the rate constancy among branches relies on the neutral condition. The substitution rate under selective neutrality depends only on the mutation rate and is independent of the population size and the selection coefficient. If the mutation rate is similar among lineages, the substitution rate can be expected to be similar among lineages. In contrast, under natural selection, the substitution rate is related to the population size and the selection coefficient. Even if a constant mutation rate is assumed, the population size and the selection coefficient are unlikely to always be constant among the lineages. Hence, rates would vary substantially among lineages, influencing the rate constancy among branches (Ohta and Kimura 1971; Takahata 1987; Ohta 1992; Nei et al. 2010; Gaut et al. 2011).

Moreover, as noted by other researchers, neutral theory also implies that the rate constancy within a branch relies on the neutral condition (Phillips and Penny 2003; Ho and Larson 2006; Subramanian et al. 2009; Subramanian and Lambert 2011). In practice, we do not distinguish whether the observed genetic variations have been fixed or not in the population; therefore, the “rate” that we refer is actually not equivalent to the substitution rate or the mutation rate (Ho et al., 2005; Subramanian and Lambert, 2012). Consider a pair of sequences. If the two sequences diverged in the very recent past, almost all the observed genetic variations are new mutations, such that the short-term rate is approximately equal to the mutation rate. However, if the two sequences diverged a long time ago, then almost all the observed genetic variations are mutations that have been fixed in the population (substitutions); thus, the long-term rate is approximately equal to the substitution rate. The “rate” undergoes a transition between the substitution rate and the mutation rate. Under selective neutrality, because the substitution rate is equal to the mutation rate, the long-term rate is equal to the short-term rate, and the “rate” is expected to be generally constant through time. Instead, under purifying selection, because the substitution rate under purifying selection is lower than the mutation rate, a phenomenon called the “time dependency of molecular rates” (TDMR) is expected: the “rate” decays as moving backward in time (Ho et al. 2005, 2015; Subramanian et al. 2009; Subramanian and Lambert 2011, 2012; Nicolaisen and Desai 2012; Ho 2014; Aiewsakun and Katzourakis 2015, 2016).

As described above, both the rate constancies among lineages and through time rely on the neutral condition. From this point of view, purifying selection — the most common type of natural selection— can be inferred as likely changing the pattern of the molecular clock, which may reduce the reliability of the result of molecular dating. In practices of species-level molecular dating, researchers have paid a great deal of attention to factors that might increase the uncertainty of the analysis, such as substitution saturation, the rate heterogeneity among sites and the uncertainty in fossil calibration (Brandley et al. 2011; Nakatani et al. 2011; Zheng et al. 2011; Soubrier et al. 2012; Zhu et al. 2015; Angelis et al. 2018). Among these factors, substitution saturation may be one of the most well-known issues. As substitution saturation could cause an underestimation of branch lengths, some researchers have proposed or adopted the selection of slowly evolving genes/sites (such as 1^st^ and 2^nd^ codon positions) to reduce the risk of being influenced by substitution saturation (Miya et al. 2010; Nakatani et al. 2011; dos Reis et al. 2012, 2014; Jarvis et al. 2014; Hu et al. 2017; Liu et al. 2017). However, from the viewpoint of purifying selection, this data processing method leads to genes/sites under neutrality being excluded and genes/sites under strong impacts of purifying selection being retained. Hence, a need exists to examine whether purifying selection has an impact on species-level molecular dating.

Here, we used 2242 protein-coding genes in 23 published mammal genomes to investigate the impact of purifying selection on species-level molecular dating. We grouped the 2242 genes were into 30 bins according to their overall selective constraints and compared the difference in branch lengths and time estimates among bins. Meanwhile, we also randomly sampled genes from the 2242 genes and compared the branch lengths and time estimates among different codon positions in these genes. Through these comparisons, we examined whether differences exist among the results of datasets under various selective constraints.

## Methods

We used the molecular dating program MCMCTree in the PAML package to perform divergence time estimation for the investigation. In the intermediate process (usedata=3), branch lengths would also be estimated by the program BaseML and written into a file named “out.BV” to facilitate the calculation of likelihood (Thorne et al. 1998; dos Reis and Yang 2011). Since the inferred branch lengths are directly related to divergence time estimation, they were used to investigate the pattern of branch lengths. If the data are partitioned, more than one phylogram tree will be present in the out.BV file, and each tree corresponds to a partition. Specifically, if the data are partitioned by codon positions, three trees corresponding to the 1^st^, 2^nd^ and 3^rd^ codon positions, respectively, will be present in the out.BV file (dos Reis and Yang 2011).

We collected 2242 coding sequences (CDS) from 23 mammalian genomes (Figure 1) and grouped them into 30 bins according to their mean pairwise d*N*/d*S* (*ω*) values. The overall selective constraint of the bin is stronger when the *ω* value is smaller. Within a bin, the selective constraint is 3^rd^ positions < 1^st^ positions < 2^nd^ positions. We evaluated the impact of purifying selection through comparisons of different datasets. To make the branches and time estimates comparable among the different datasets, the following described analyses were performed with the same topology (see the topology in Figure 1). The comparisons among the bins were performed under 5 different schemes: using only 1^st^ positions, using only 2^nd^ positions, using only 3^rd^ positions, using all sites of genes under concatenation and using all sites of genes under partitioning by codon positions. Meanwhile, we also compared different codon positions in randomly sampled genes (see an illustration in Figure 2).

**Figure 1.**
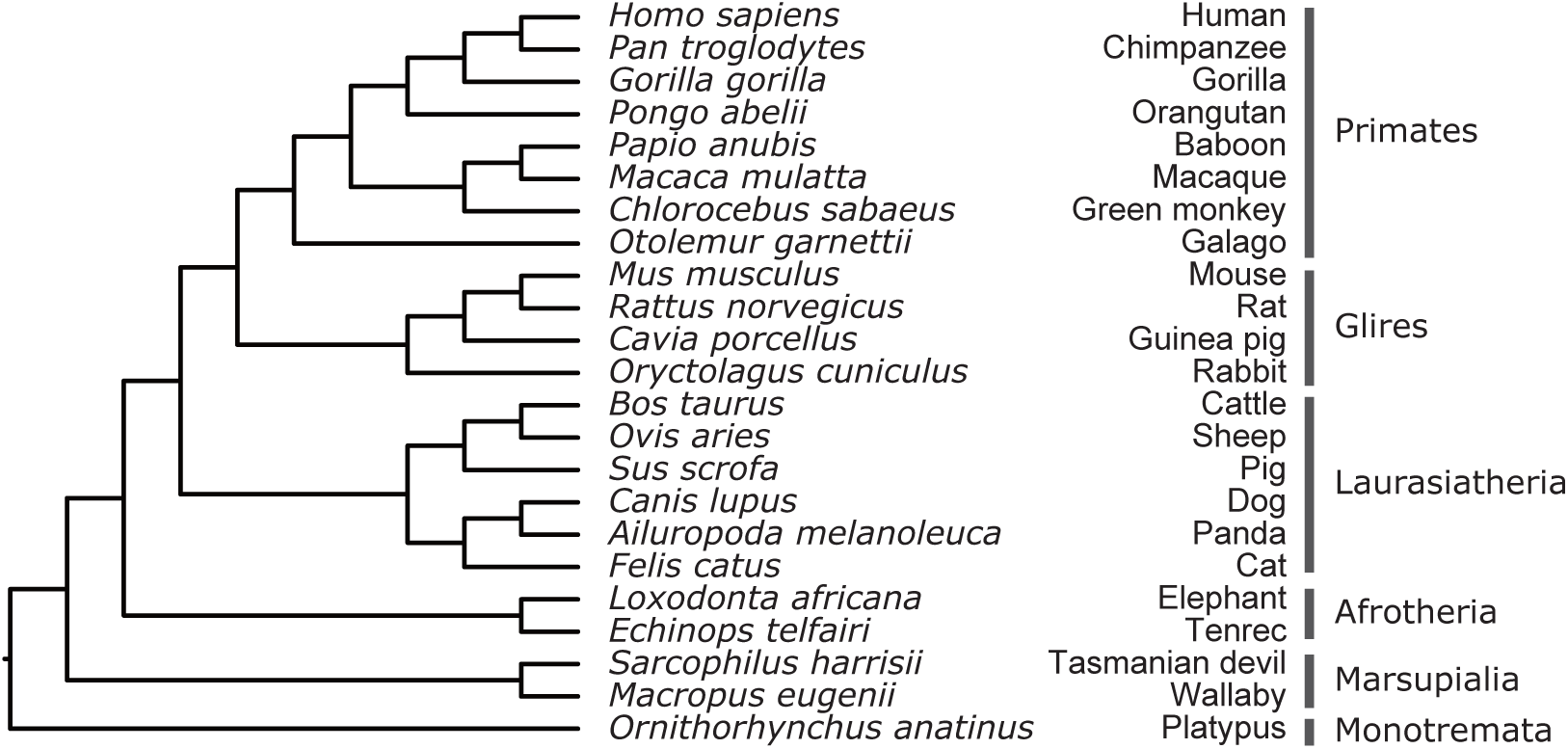
The 23 mammals and topology used for investigating the impact of purifying selection on species-level molecular dating.

**Figure 2.**
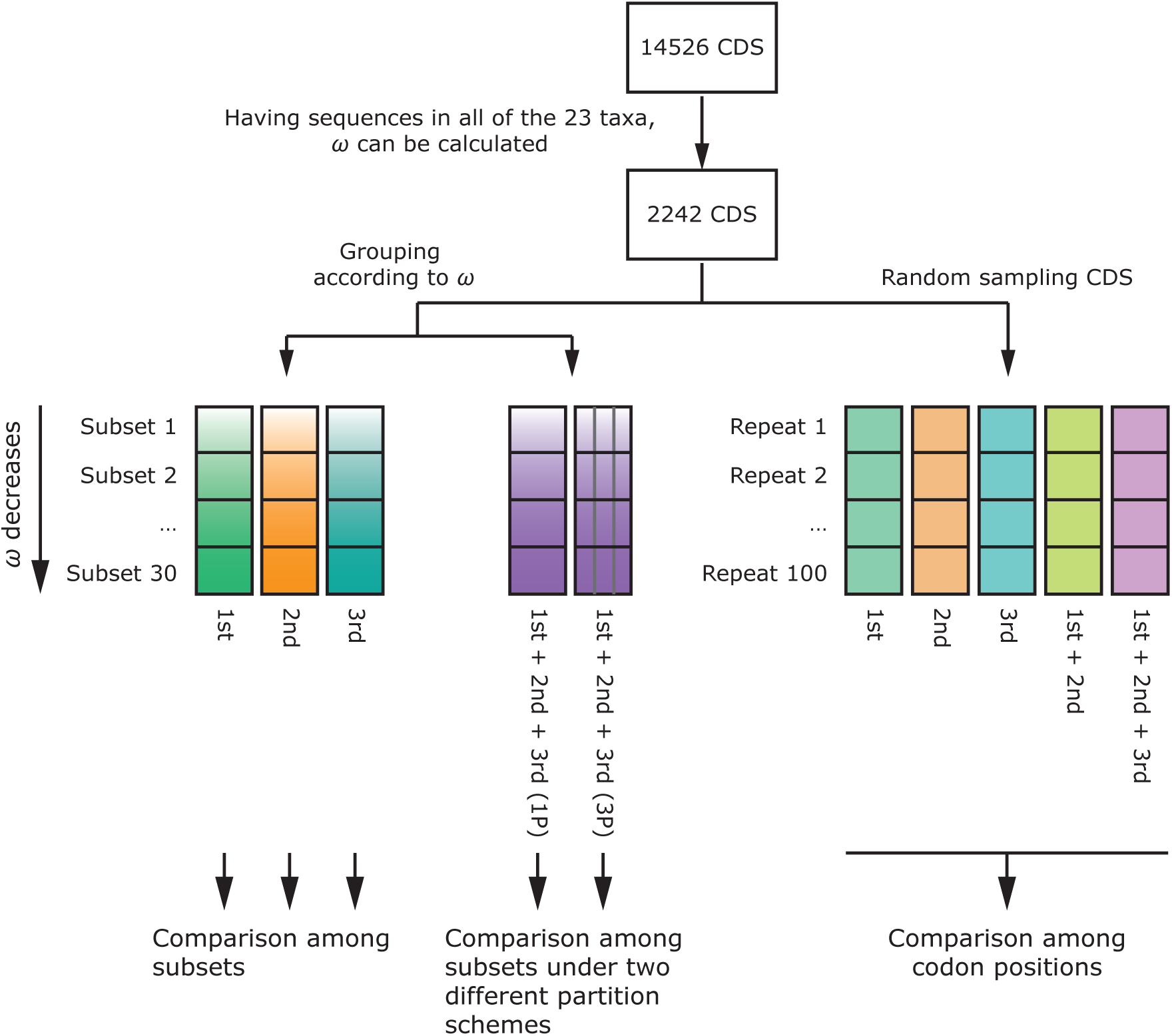
The workflow of investigating the impact of purifying selection on species-level molecular dating.

### Obtaining and Filtering Coding Sequences (CDS)

We selected 23 mammal species that represent major mammalian lineages for our study. Based on previous studies, the divergence times among these species range from 5 Ma to 185 Ma (Meredith et al. 2011; dos Reis et al. 2012, 2014). A total of 14,526 mammal CDS alignments were downloaded from the OrthoMaM database (Douzery et al. 2014). To minimize the influence of missing data, we chose the CDS alignments that have sequences of all the selected taxa for further analyses. The reason why we selected twenty-three rather than all available mammal species is to obtain more genes that satisfy the above criteria. Mitochondrial protein-coding genes were discarded. Mean pairwise d*N*/d*S* (*ω*) was used to measure the overall selective constraint on a CDS. To calculate *ω*, pairwise nonsynonymous substitutions (d*N*) and synonymous substitutions (d*S*) were calculated by the CodeML program in the PAML package (Yang 2007), and *ω* was calculated as (mean d*N*)/(mean d*S*). For some CDS, *ω* cannot be calculated because no site was retained or because no difference existed in the retained sites after deleting the gaps; thus, they were excluded from analyses. Finally, 2242 CDS alignments were retained for further analyses.

### Workflow of the Investigation

We analyzed both relative branch lengths and the time estimates under different selective constraints. The workflow of the investigation is shown in Figure 2. The 2242 CDS were ranked by *ω* and grouped into 30 bins. When *ω* is small (under strong selective constraint), the variable sites in the 2^nd^ positions may not be sufficient to precisely estimate branch lengths and divergence times; thus, in the grouping procedure, we made 30 bins with similar numbers of variable sites in the 2^nd^ codon positions rather than making these bins with similar numbers of genes or informative sites.

First, we considered the effects of the gene and the codon position. We separated the 1^st^, 2^nd^, and 3^rd^ codon positions to perform the investigation. For each of the 30 bins, three phylogram trees and time trees were estimated based on the different codon positions. Correspondingly, three linear regressions were performed to detect the impact of purifying selection. Note that, although linear regressions were performed here, we did not suggest any linear relationship between *x* and *y*; it was only used to detect whether a systematic impact exists. The *p*-value of the linear regression indicates the probability that the slope is zero, i.e., the probability that the datasets fluctuate randomly around a constant value. Thus, if the *p*-value is significantly small, it indicates the existence of a systematic impact.

Next, we combined all the codon positions together to compare the overall difference among bins. Considering the impact of partitioning scheme, the investigations were conducted under two different partitioning schemes: concatenating all sites as one partition (1P) and partitioning by codon position (3P). As mentioned in the beginning of the Methods, the time tree under the 3P scheme is based on the three phylogram trees (also the gradient vectors and Hessian matrix) that correspond to the three codon positions. These phylogram trees are same as what we investigated above. Thus, the pattern of the branch lengths for the 3P scheme is exactly the same as what we investigated above, and no need exists to perform the same investigation. To summarize, for each of the 30 bins, one phylogram tree under the 1P partitioning scheme and two time trees corresponding to the two partitioning schemes were to be estimated in this part of investigation. Accordingly, only one linear regression was performed to detect the impact on the branch length, and two were performed to detect the impact on the time estimate.

Next, we randomly sampled 100 CDS from the 2242 CDS with 100 repetitions, and we investigated the behaviors of the different codon positions. For each repeat, we conducted five different treatments: using the 1^st^ codon position, 2^nd^ codon position, 3^rd^ codon position, 1^st^ + 2^nd^ codon positions and 1^st^ + 2^nd^ + 3^rd^ codon positions. We compared the differences among these treatments. Correspondingly, in each repeat, 5 phylogram trees and time trees have to be estimated and compared.

We wrote Python scripts to implement these procedures. Alignments and tree files were parsed by Biopython to facilitate extracting sequences and branch length information (Cock et al. 2009; Talevich et al. 2012). Linear regressions were performed by the SciPy library to calculate regression equations and *p*-values (Millman and Aivazis 2011). Plots were drawn by matplotlib library (Hunter 2007). Details of the aforementioned procedures are described in the following sections.

### Estimation of Branch Lengths and Divergence Times

The program MCMCTree in the PAML package was used in the present study (Yang and Rannala 2006; Rannala and Yang 2007; Yang 2007; dos Reis and Yang 2011). We used the approximate likelihood method (dos Reis and Yang 2011) following a step-by-step protocol written by the developers running the program. The gradient vector, Hessian matrix and branch lengths were inferred under the HKY85 + *Γ*_4_ by the program BaseML (Yang 2007) with reference to a previous study, dos Reis et al. (2014). For all the datasets, the tree shown in Figure 1 was used as the reference topology. As mentioned above, the inferred branch lengths in this step were used to investigate the pattern of branch lengths. We additionally ran phylogenetic reconstruction program RAxML (Stamatakis 2014) without fixing topology to examine whether the result is an artefact caused by the mismatch between topology and data.

The divergence times were estimated in MCMCTree with setting “usedata” as 2 under the auto-correlated rate model (1,000,000 iterations; first 10% as burn-in). The shape parameter of gamma prior for the overall rates for genes (“rgene_gamma”) was set as 2, and the gamma prior for rate drift (“sigma2_gamma”) was set as G(1, 1). Divergence time estimations were run at least twice to test whether the MCMC had reached convergence. Time estimates among bins are comparable only if they have a “common starting point”. Note that under a reversible substitution model (e.g. HKY85, GTR), there is no way to know the distance between the root of the whole tree (the crown Mammalia) and the second basal node (the crown Theria) just based on the molecular data (i.e. Felsenstein’s “pulley principle”) (Felsenstein 1981). If we calibrate only the root of the tree, the time of the second basal node can be varied among datasets. However, such a variation is irrelevant to the factor that we are interested in (the relative branch length). Therefore, to set a “common starting point”, the second basal node (or in another word, the root of the in-group) needs to be calibrated (similar rationale can be seen in Thorne et al., 1998). We calibrated the root and the second basal node with tightly constraints >1.8579<1.8581 and >1.7019<1.7021. They were according to the estimated divergence times of (dos Reis et al. (2014). This calibration scheme forces the time estimates of the root and the second basal node to be nearly identical among datasets, thus providing a “common starting point”. Under this calibration scheme the time estimates of the other 20 nodes are comparable; and we did not calibrate any other node, thus the influence of the change in relative branch lengths can be shown in the maximum extent.

### Measures of the Relative Branch Length and Branch Variation

We used the ratio of the sum of the terminal branch lengths to the sum of the internal branch lengths (*SumT*/*SumI*) to measure the overall relative length of terminal branches,

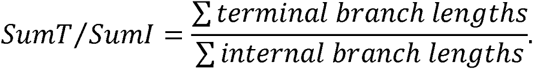

The ratio of each terminal branch length to the sum of internal branch lengths (*T*/*SumI*) was used to measure the relative length of each terminal branch,

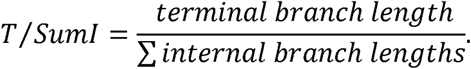

We used the coefficient of variation (CV) of node-to-tip distances to study the impact on rate heterogeneity.

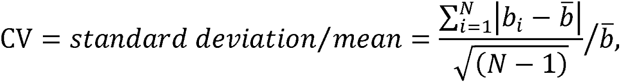

where *N* is the number of lineages, *b_i_* is the distance from the tip of *i*-th lineage to the node of the most recent common ancestor (MRCA) of the *N* lineages, 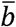 is the mean of node-to-tip distances, 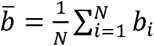.

## Results and Discussion

### The Change in the Branch Length as the Selective Constraint Becomes Stronger

In species-level molecular dating, the role of sequence data is to provide information about genetic distances (branch lengths) (dos Reis et al. 2016). Therefore, we first show the pattern of the relative branch lengths among the different bins. To visually display our observations, the branch lengths inferred from three representative bins are given in Figure 3: (1) bin #1, under the most relaxed selective constraint (*ω* = 0.48); (2) bin #17, under a moderate selective constraint (*ω* = 0.12); and (3) bin #30, under the most rigid selective constraint (*ω* = 0.01). Let us start with the 3^rd^ positions of bin #1, which is under the most relaxed selective constraint. We use an indicative node, Catarrhini (including human, chimpanzee, gorilla, orangutan, baboon, macaque and green monkey), to help us clarify our observation. For the 3^rd^ positions of bin #1, the node-to-tip distances for Catarrhini were similar, showing relatively constant rates for this group. Additionally, for all the codon positions of bin #1 and for 3^rd^ codon positions among the three representative bins, the shapes of the trees were similar (Figure 3). This pattern is consistent with the rate constancy under the neutral condition, which has been highlighted by a series of early studies. As the selective constraint becomes stronger, the shapes of the trees became distorted. As one of the signatures of the distortion, the variation among the node-to-tip distances for crown Catarrhini became increasingly large (from the lower left to the upper right in Figure 3). To show the observation more quantitatively, we performed linear regressions for the three kinds of codon positions with the coefficient of variation (CV) of node-to-tip distances for crown Catarrhini as the scalar response (*y*) and the *ω* of the corresponding dataset as the explanatory variable (*x*). For the 3^rd^ positions, the CV was quite similar across bins; however, for the 1^st^ and 2^nd^ positions, we found that as *ω* decreased, the CV increases (slope > 0), and the trend of the 2^nd^ positions has a larger slope value than that of the 1^st^ positions (Figure S1). This pattern seems to be consistent with the idea that the existence of natural selection can increase the rate heterogeneity among the lineages (Ohta and Kimura 1971; Ohta 1992; Gaut et al. 1996).

**Figure 3.**
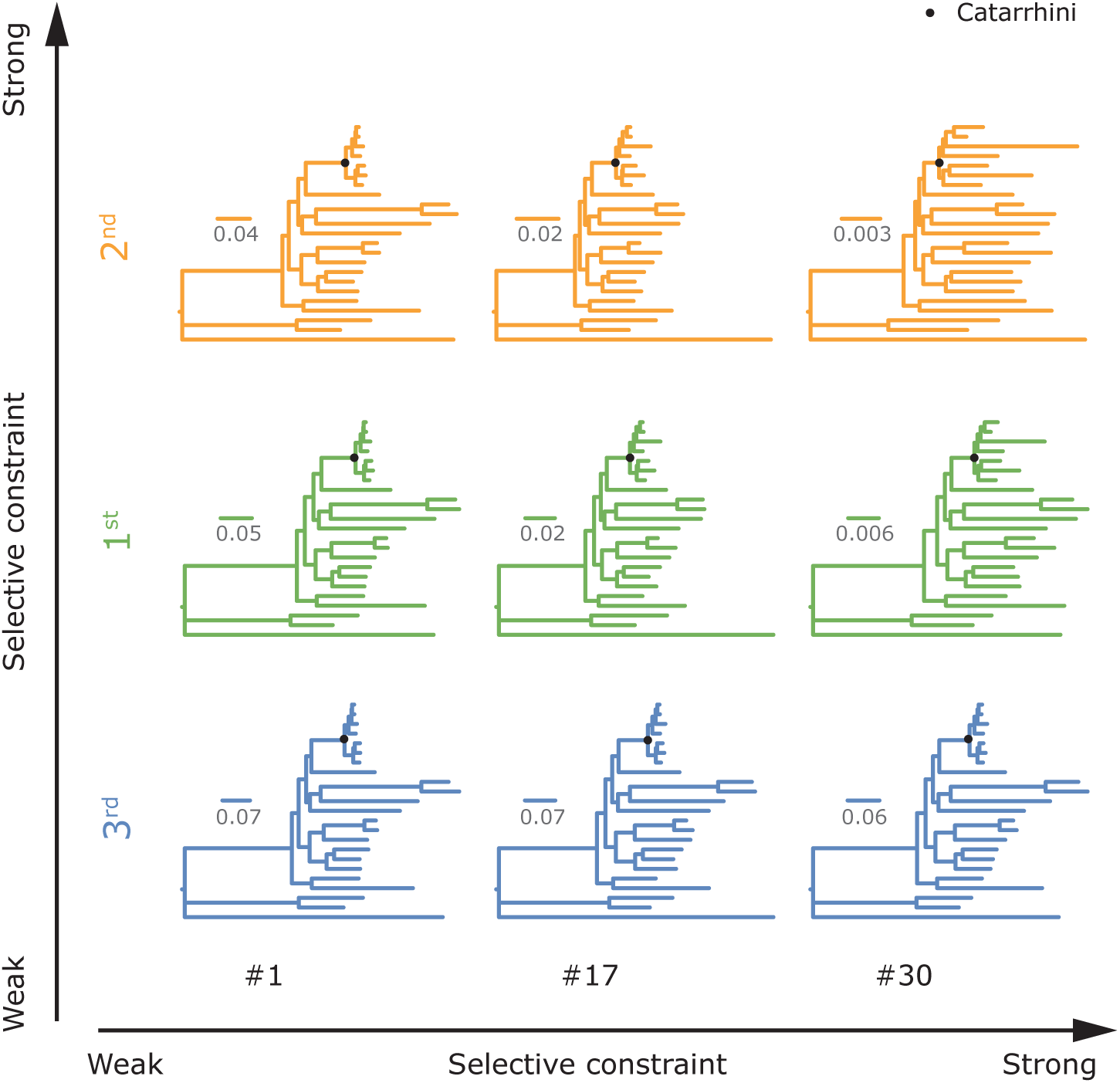
The branch lengths of three representative bins. The branch lengths shown here were inferred from different codon positons of three representative bins “#1”, “#17” and “#30”, which are under the least, moderate and strongest selective constraints, respectively. The topology of each tree follows that of Fig. 2.

The distortions of trees did not just show a pattern in which the branches of some lineages were lessened and those of others were extended. Instead, we noted that, as the selective constraint became stronger, almost all the terminal branches became relatively extended (they were lessened in terms of the absolute value). For each lineage, we performed linear regressions with the ratio of the length of each terminal branch to the sum of all internal branch lengths (*T*/*SumI*) as the scalar response (*y*) and *ω* as the explanatory variable (*x*). We found that for the 1^st^ and 2^nd^ codon positions, all the trends had positive slopes (Figure 4; with exceptions that *p* > 0.05 for both 1^st^ and 2^nd^ positions in mouse, and for 1^st^ positions in chimpanzee and wallaby). The existence of such a large proportion of terminal branches showing positive slope values in the linear regressions is statistically significant (see Supplementary Methods and Table S1). Hence, the observed extension of the terminal branches is unlikely due to chance or lineage-specific adaptations. Additionally, we performed linear regressions with the ratio of the sum of terminal branch lengths to the sum of internal branch lengths (*SumT*/*SumI*) as the scalar response (*y*) and *ω* as the explanatory variable (*x*). For 3^rd^ positions, *SumT*/*SumI* values were generally similar among the 30 bins. For both 1^st^ and 2^nd^ codon positions, *SumT*/*SumI* values increased significantly as *ω* decreased (slope > 0, *p* < 0.05), and the trend for the 2^nd^ positions has a larger slope value than that of the 1^st^ positions (Figure 4). This pattern remained stable when trees were estimated by the phylogenetic reconstruction program RAxML without fixing the topology (Figure S2). Thus, the extension of the terminal branches is also unlikely to be due to the mismatch between the topology and data.

**Figure 4.**
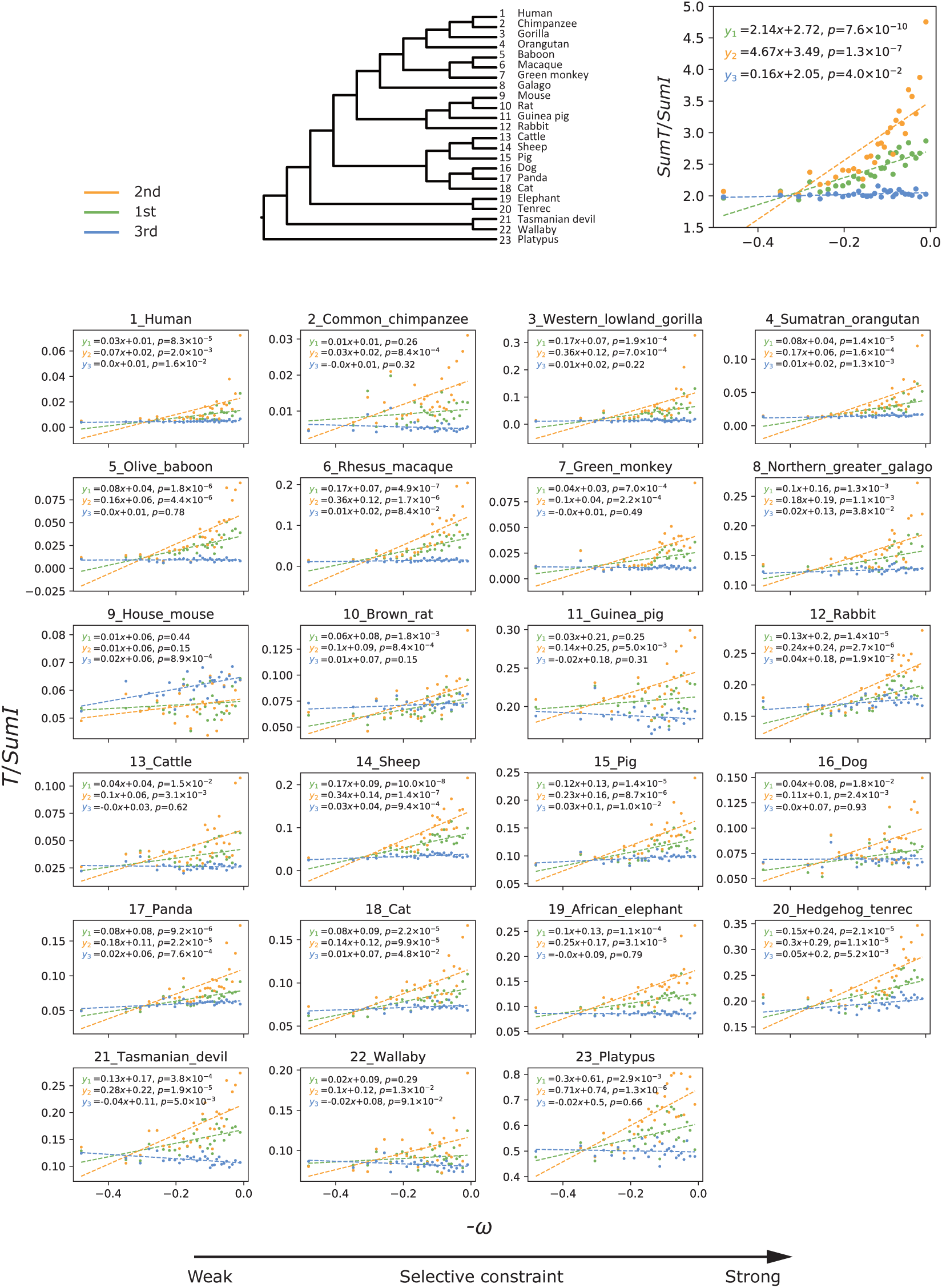
The change in the branch length as the selective constraint becomes stronger. The *x*-axis is the opposite of the mean pairwise d*N/*d*S* (-*ω*), indicating the overall selective constraint on a bin (the right is under the stronger constraint). At the upper, the *y*-axis is the ratio of the sum of terminal branch lengths to the sum of internal branch lengths (*SumT*/*SumI*), indicating the overall relative length of terminal branches. At the lower, the *y*-axis is the ratio of the branch length of each terminal branch to the sum of internal branch lengths (*T*/*SumI*), indicating the relative length of each terminal branch. Overall, as the selective constraint becomes stronger, the terminal branches are relatively extended. Such a change in the branch length can be detected for almost all the terminal branches.

### The Change in the Time Estimate as the Selective Constraint Becomes Stronger

Next, we show the pattern of time estimates. Time estimates among datasets can be comparable only if they share a “common starting point”. We calibrated the root of the in-group with tight constraints (based on the result of a previous study) (dos Reis et al., 2014) to force the time estimate for this node to be nearly identical among datasets, thus providing a “common starting point” (see Methods). Under this calibration scheme, the divergence times of the other 20 nodes were estimated and compared (note that the branch length estimation is independent of the calibration scheme; regardless of which calibration scheme is adopted, the above pattern of branch lengths holds).

The most marked effect on the time estimate is correlated with the extension of the terminal branches. Overall, the time estimates based on the 1^st^ and 2^nd^ codon positions become older as *ω* decreased, and the trends for the 2^nd^ codon positions had larger slope values than those for the 1^st^ codon positions; whereas, for the 3^rd^ positions, the time estimates were similar among the different bins (Figure 5; see representative time trees in Figure S3). For the 2^nd^ codon positions, all the nodes showed regression trends with positive slope values (*p* < 0.05 in binominal test, see Supplementary Methods and Table S2), 16 of which showed statistical significances; and the other 4 nodes that did not show statistical significance were older than 90 Ma. For the 1^st^ codon position, 18 of the 20 nodes showed regression trends with positive slope values (*p* < 0.05 in binominal test, see Supplementary Methods and Table S2), 11 of which showed statistical significances; the other 9 nodes that did not show statistical significance were older than 80 Ma.

**Figure 5.**
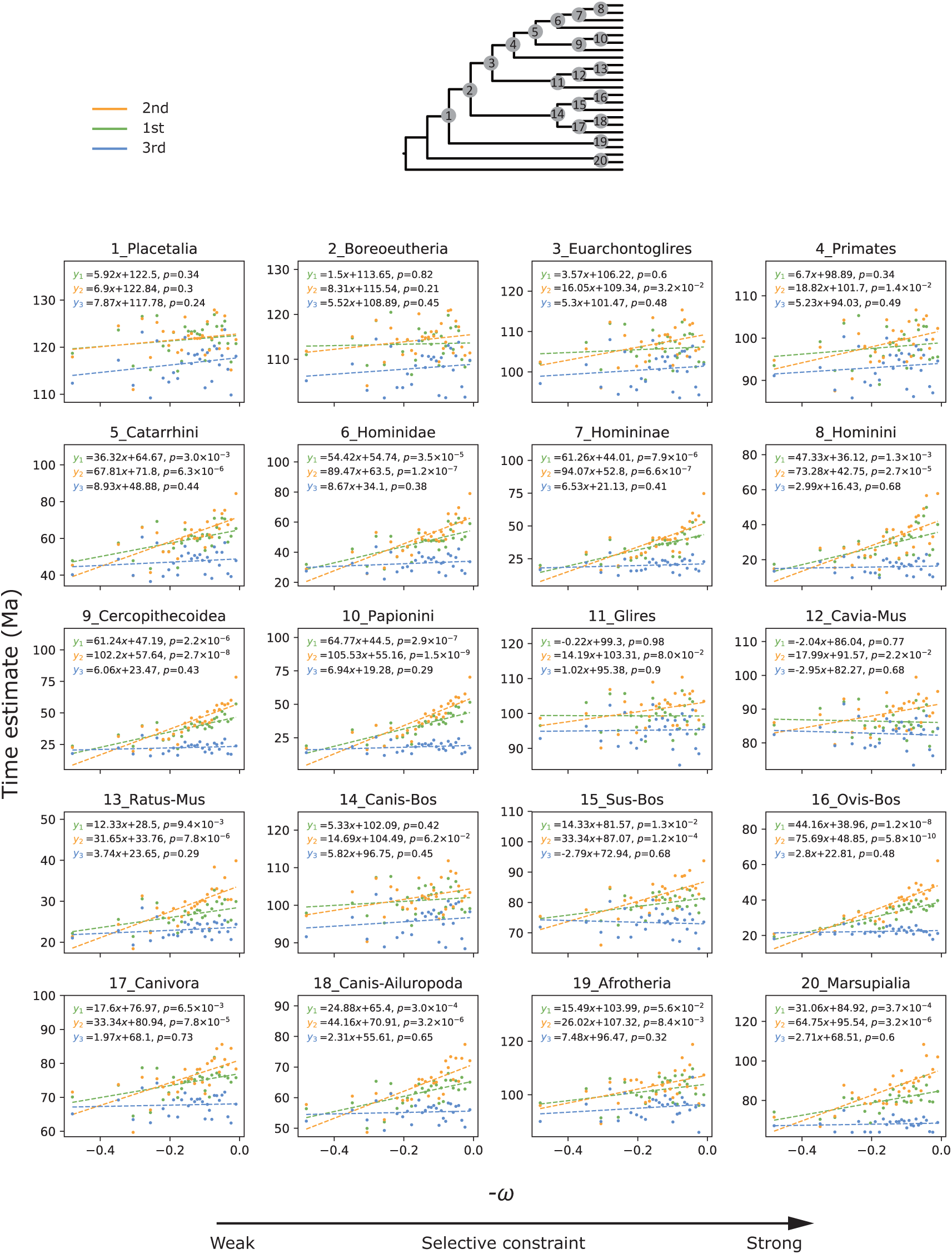
The change in the time estimate as the selective constraint becomes stronger. The *x*-axis is the opposite of the mean pairwise d*N/*d*S* (-*ω*), indicating the overall selective constraint of a bin (the right is under the stronger constraint). The *y*-axis is the time estimate for each node. Overall, as the selective constraint becomes more rigid, the time estimates become older. The shallow-scale nodes are impacted more severely than deep nodes.

The impact on the time estimate was more pronounced for shallow-scale nodes than deep-scale nodes (Figure S4). For example, for crown Primates (node 4, Figure 5; a deep-scale node), the time estimate of the 2^nd^ positions in bin #30 was 12.27% older than of the 3^rd^ positions in bin #1 (102.29 Ma vs. 91.11 Ma), while, for the crown Papionini (node 10, Figure 5; a shallow-scale node), the time estimate of the 2^nd^ position in bin #30 was 407% older than that of the 3^rd^ position in bin #1 (70.28 Ma vs. 13.86 Ma). These results, combined with the above results for branch lengths, show that the extended terminal branches can “push” the time estimates to be older as the selective constraint becomes stronger. Accordingly, purifying selection can influence the result of species-level molecular dating.

### The Change in the Branch Length and the Time Estimate When Using All Sites of Genes

In the above analyses, the three codon positions were separated for each bin. It is also worth investigating the overall behaviors of bins using all the three codon positions of genes. Here, we compared the 30 bins with using all the three codon positions together. As different codon positions are involved, a consideration of the impact of partitioning scheme is required. Thus, we conducted the comparison of time estimates under two treatments: concatenating all sites as one partition and partitioning the data into three partitions according to codon positions (see Methods and Figure 2). Note that with partitioning by codon positions, the time tree is based on the branch lengths of the three phylogram trees that correspond to the three codon positions (see Methods). For these trees, we have already analyzed and discussed above. In this part of investigation there is no need to discuss this result again, thus the investigation of branch lengths was performed only for the 1P scheme.

Let us start with the result for the 1P scheme, where each bin corresponds to a single phylogram tree and the time tree is based on this tree. We found that when all sites were concatenated as one partition, *SumT*/*SumI* values of bins also showed an increasing trend as *ω* decreased, but the slope value was small (Figure 6, upper), suggesting a modest impact of purifying selection. Consistent with the pattern of branch lengths, time estimates under 1P scheme also showed some increases as *ω* decreased (Figure 6). For 19 out of the 20 nodes, the slope values were positive (*p* < 0.05 in binominal test, see Supplementary Methods and Table S2). Nevertheless, the difference in time estimates among bins were modest (see representative time trees in Figure S5). The regression trends had smaller slope values than the trends for 1^st^ and 2^nd^ codon positions and only 7 nodes showed statistical significances (Figure 6). With a consideration of the neutral theory, this result seems to be not surprising. As suggested by the neutral theory, in general, most of the observed genetic variations are selectively neutral (Kimura 1968, 1977; Ohta 1992; Nei et al. 2010). Without artificial manipulation, neutral substitutions (majorly from 3^rd^ positions) are expected to be the major contributors for the branch length. Hence, the overall behavior of a gene should be similar to that of its 3^rd^ positions, differences among bins would not be substantial.

**Figure 6.**
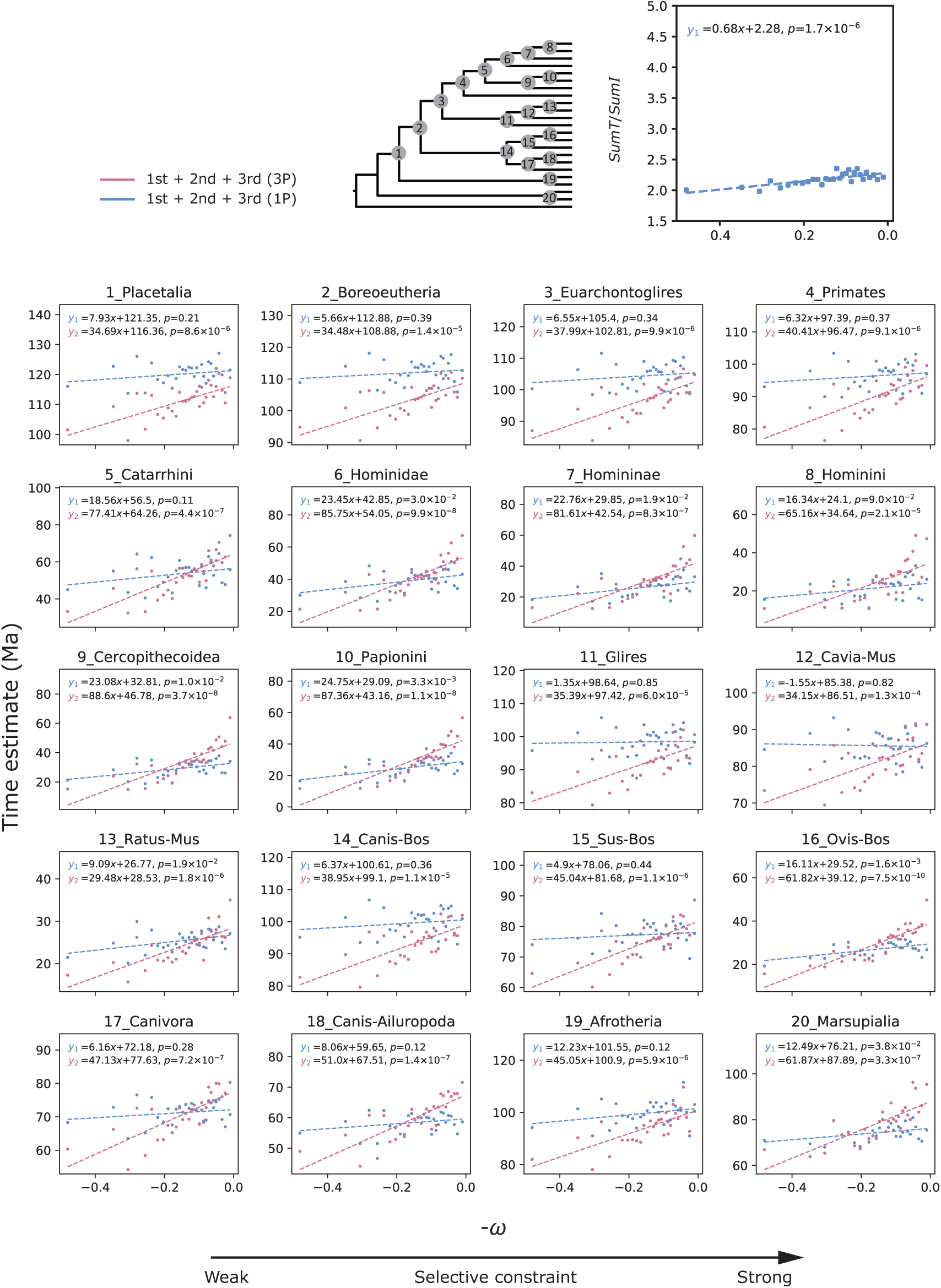
The change in the branch length and the time estimate when using all sites of genes. The patterns under two different partitioning schemes, concatenating all sites as one partition (1P) and partitioning by codon position (3P) are shown. The *x*-axis is the opposite of the mean pairwise d*N/*d*S* (-*ω*), indicating the overall selective constraint of a bin (the right is under the stronger constraint). At the upper, the *y*-axis is the ratio of the sum of terminal branch lengths to the sum of internal branch lengths (*SumT*/*SumI*) based on all sites of genes. The linear regression shows a positive slope, however, it is the slope value is small, which suggests that when using all sites of genes, although the extension of the terminal branches can be detected, the extent is modest. At the lower, the *y*-axis is the time estimate for each node. Under the 1P scheme (blue), the difference in time estimates among bins were not prominent; the slope values of the linear regressions are generally small. However, under the 3P scheme (purple), the difference in time estimates among bins become prominent; the slope values of the linear regressions are much larger than under the 1P scheme. The impact of purifying selection on the time estimate under the 3P scheme is stronger than under the 1P scheme.

Nevertheless, under the 3P scheme the pattern became different. We found that under the 3P scheme, as *ω* decreased the time estimates showed much more prominent increases than under 1P scheme (see representative time trees in Figure S5). The regression trends had larger slope values than the trends under 1P scheme and all the regression trends showed positive slope values and had statistical significances (Figure 6, Table S2 and Supplementary Methods). The pattern under 3P scheme is more similar to that of 1^st^ and 2^nd^ positions rather than that of 3^rd^ positions. The mechanism behind this result could be complicated. But one thing should be noted here: in the algorithm of molecular dating, the divergence times of different partitions are assumed to fluctuate up and down randomly around a “true tree” (Thorne and Kishino 2002; Yang and Rannala 2006; dos Reis and Yang 2011). According to the above results, this assumption is violated under purifying selection. The impact of purifying selection may thus be strengthened.

In summary, when all sites of genes are used together, the impact of purifying selection can also be detectable. The strength of the impact of purifying selection depends on the partition scheme. Under concatenating all sites as one partition, the differences among bins are small, the impact of purifying selection is generally modest. While, under partitioning by codon position, the differences among bins become substantial, the impact of purifying selection is strengthened. Rate heterogeneity among codon positions is usually larger than that among genes. Some researchers would partition the data by codon position to accommodate such rate heterogeneity (Yang and Rannala 2006; Brandley et al. 2011; Shen et al. 2016; Liu et al. 2017; Angelis et al. 2018; Morris et al. 2018). Nevertheless, considering the impact of purifying selection, this partitioning strategy could be problematic. We suggest researchers being more cautious about this method in future.

### The Result of the Comparison among Different Codon Positions in Randomly Sampled Genes

In species-level molecular dating practices, the removal of the 3^rd^ codon positions and use only the 1^st^ and 2^nd^ codon positions are common to avoid the potential impact of substitution saturation. However, the sites at the 1^st^ and 2^nd^ codon positions are typically under stronger purifying selection. To evaluate the influence of such a practice, we generated 100 randomly sampled datasets, each of which contained 100 CDS from the 2242 CDS. For each dataset, we estimated the branch lengths and divergence times by using only the 1^st^ codon positions, only the 2^nd^ codon positions, only the 3^rd^ codon positions, 1^st^ + 2^nd^ positions and all sites. In all the 100 randomly sampled datasets, the *SumT*/*SumI* values were as follows: the 2^nd^ position > 1^st^ + 2^nd^ positions > 1^st^ position > all sites > 3^rd^ position (Figure 7, upper), and all pairwise comparisons showed statistical significance (Supplementary Methods, Table S2). Correspondingly, the mean time estimates of the 20 nodes were as follows: the 2^nd^ position > 1^st^ + 2^nd^ positions > 1^st^ position > all sites > 3^rd^ position. The time estimates based on the 3^rd^ position were consistently the youngest, the time estimates were older under the stronger selective constraint of the dataset (Figure 7), and all the pairwise comparisons showed statistical significance (see Supplementary Methods, Table S3). Specifically, for the widely adopted practice of using 1^st^ + 2^nd^ positions, nodes not older than 40 Ma could produce ∼ 20% to 50% older time estimates than those determined by using all sites. Hence, for practices such as using the 1^st^ + 2^nd^ positions, the impact of purifying selection should not be neglected.

**Figure 7.**
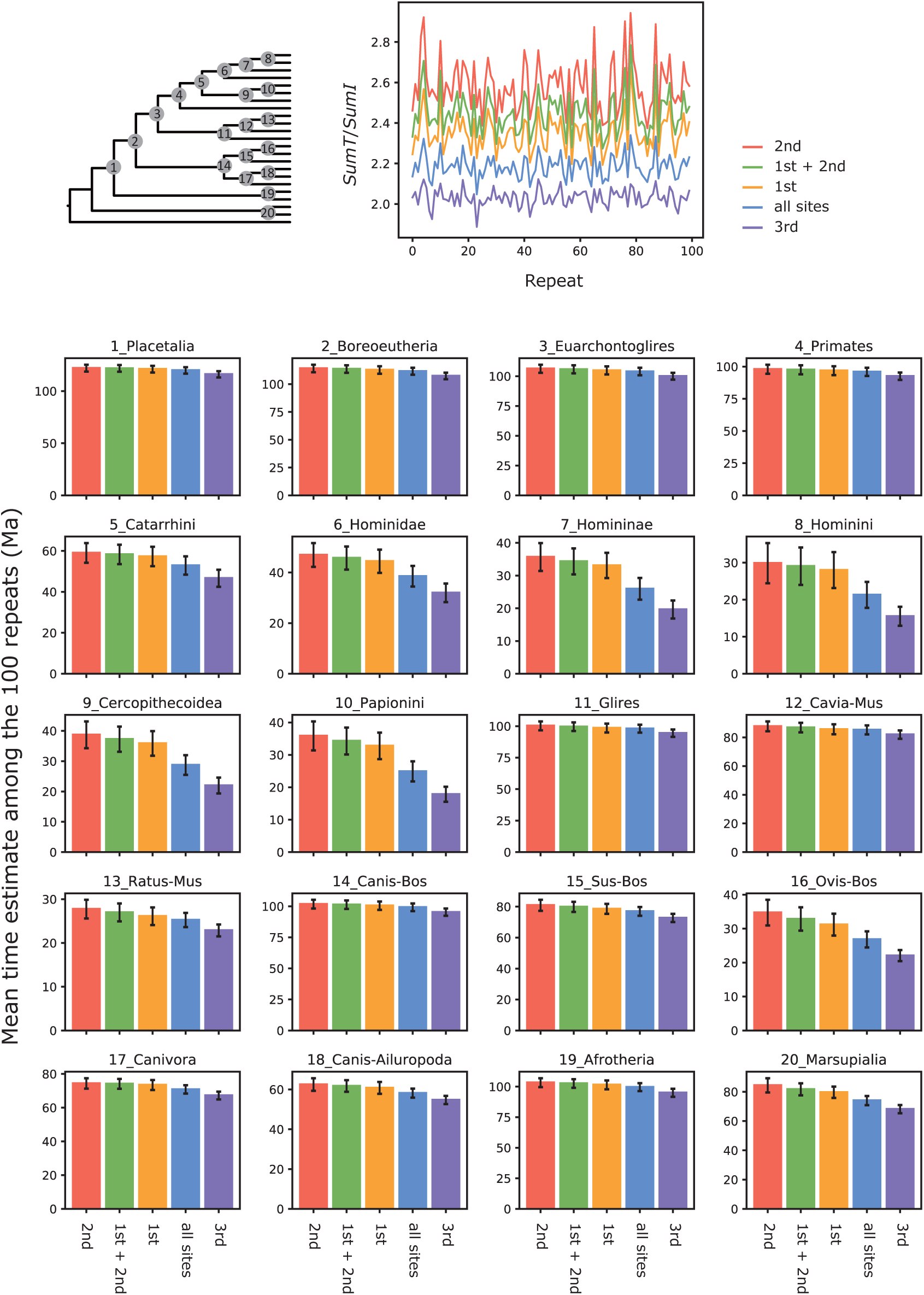
The comparison among different codon positions in randomly sampled genes. The upper panel shows the ratio of the sum of terminal branch lengths to the sum of internal branch lengths (*SumT*/*SumI*) of the 1^st^ position, 2^nd^ position, 3^rd^ position, 1^st^ + 2^nd^ positions and all sites for the 100 randomly sampled repeats (each of which includes 100 genes). In general, the *SumT*/*SumI* values are ranked as the 2^nd^position > 1^st^ + 2^nd^ positions > 1^st^ position > all sites > 3^rd^ position. The lower panel shows the mean time estimates of different codon positions for each node, which are also ranked as the 2^nd^ position > 1^st^ + 2^nd^ positions > 1^st^ position > all sites > 3^rd^ position.

### The Possible Cause of the Extension of the Terminal Branches

Finding an explanation for the extension of the terminal branches is helpful to better understand the impact of purifying selection. In species-level molecular dating, researchers generally equate the “rate” with the substitution rate. The substitution rate depends on the mutation rate, population size and selection coefficient. With this perspective of thinking, only if one of the above factors undergoes a kind of consistent change in all terminal branches, and such a kind of change depends on the selective constraint, the observed pattern could be expected. This situation is unlikely to happen. Thus, a change to this way of thinking is necessary.

By acknowledging that the “rate” is not equivalent to the substitution rate, the extension of the terminal branches can be explained naturally. Recall that the TDMR caused by purifying selection mentioned in Introduction, where the “rate” under purifying selection undergoes a transition from the mutation rate to the lower substitution rate moving backward in time. Moving forward in time, the TDMR caused by purifying selection is equivalent to a rate elevation. When mapped to a tree, this rate elevation extends terminal branches relative to the internal branches (Figure 8). When a certain node is calibrated, the extended terminal branches would “push” the time estimates of its descendant nodes to be older (Phillips, 2009). As the selective constraint becomes stronger, the substitution rate is increasingly reduced, while, the mutation rate is generally unaffected. Thus, the disparity between the substitution rate and the mutation rate increases, and the rate elevation is more severe. Therefore, as the selective constraint becomes stronger, the extension of the terminal branches strengthens more severely, and the overestimation of the time estimates also worsens, as we have seen in the above results (Figure 8).

**Figure 8.**
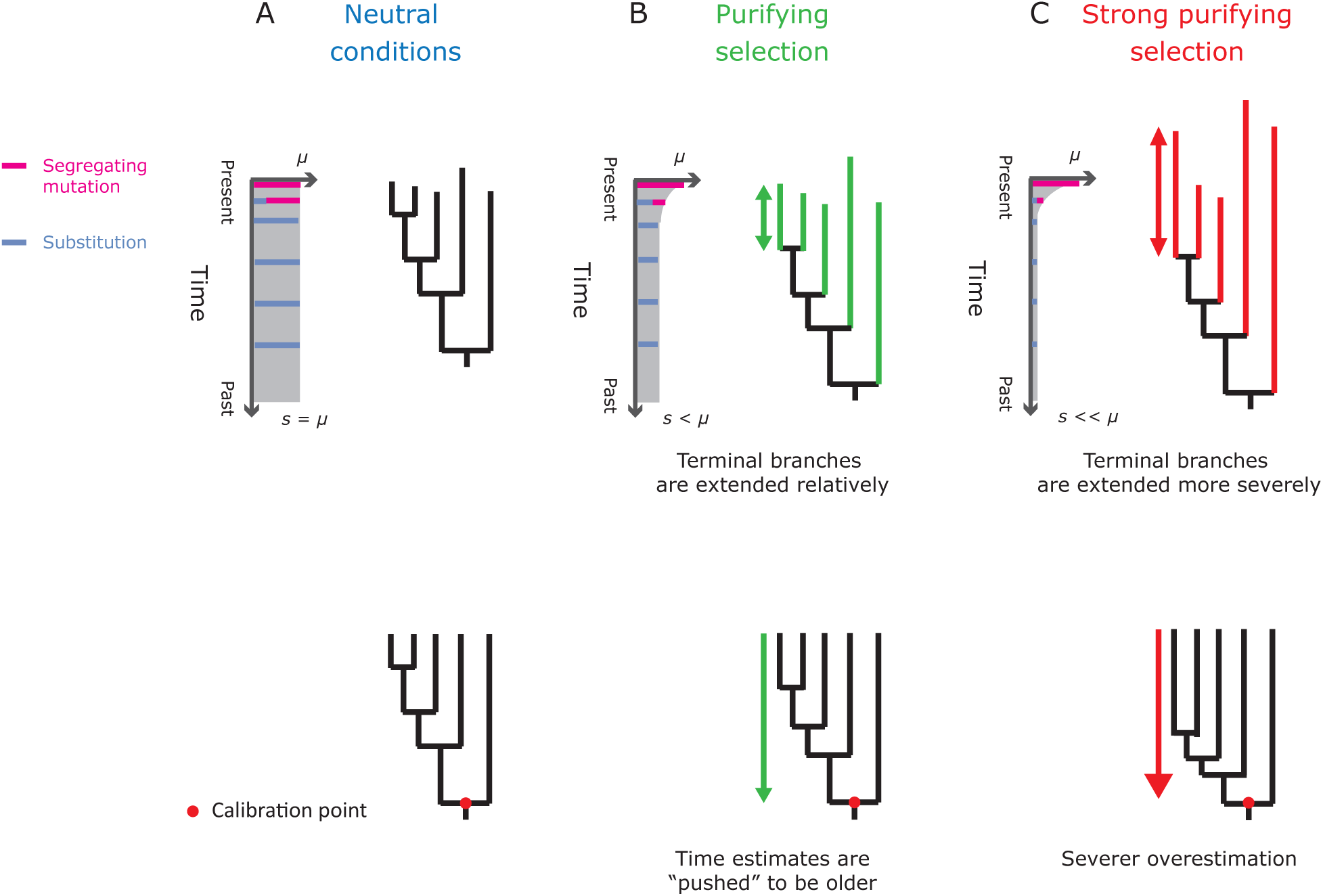
The expected effect of the time-dependency of molecular rates caused by purifying selection on branch lengths. Along the terminal branch, the “rate” undergoes a transition between the mutation rate (*μ*) and the substitution rate (*s*). A. Under neutral conditions, *s* = *μ*, the “rate” is uniform through time. B. In contrast, under purifying selection, *s* < *μ*, the “rate” elevates along the terminal branch. In this case, the terminal branch would be extended relatively. When the time of a node is calibrated, the extended terminal branches could “push” the time estimates of its descendants to be older. C. As the selective constraint becomes stronger, the substitution rate becomes smaller, thus the extension of the terminal branches becomes more severe, leading to more serious overestimation.

Can other factors lead to the extension of the terminal branches? First, we consider factors other than purifying selection that have been proposed to explain the TDMR pattern (Ho et al. 2005; Soubrier et al. 2012; dos Reis and Yang 2013). Note that, being able to explain the TDMR pattern does not directly mean being able to explain the extension of the terminal branches. Substitution saturation is one of factors that have been proposed to explain the TDMR. Substitution saturation can lead to an underestimation of branch lengths. As the distance between the sequences grows, substitution saturation tends to be more severe; thus, as the distance between the sequences grows, underestimation of branch lengths becomes more severe leading to the TDMR pattern (Ho et al. 2005, 2011). Now, let us consider if it can explain the extension of the terminal branches. Fast evolving genes are more easily influenced by substitution saturation than slowly evolving genes, as the fast evolving genes/sites are more divergent than slowly evolving genes/sites. Hence, from the viewpoint of substitution saturation, *SumI* is expected to be underestimated most seriously for the fastest-evolving dataset; the fastest-evolving dataset has the largest *SumT*/*SumI* value, and the slowest-evolving dataset has the smallest *SumT*/*SumI* value. However, the pattern that we observed in reality is opposite of this situation: the fastest-evolving dataset (3^rd^ positions of bin #1) had the smallest *SumT*/*SumI* value, and the slowest-evolving dataset (2^nd^ positions of bin #30) had the largest *SumT*/*SumI* value. Moreover, when using Xia’s tests (Xia et al. 2003), we could not detect a significant impact of substitution saturation, even for the fastest-evolving dataset (Table S4). Therefore, substitution saturation is unlikely to be the cause behind the extension of the terminal branches.

With a similar rationale, we can exclude other factors, such as selection heterogeneity among sites(dos Reis and Yang 2013) and rate heterogeneity among sites (Soubrier et al. 2012). Similar to substitution saturation, these factors can also lead to underestimation of the branch lengths. As the underestimation of branch lengths is more serious for distantly divergent sequences, a TDMR pattern can be expected (Soubrier et al. 2012; dos Reis and Yang 2013). Again, fast evolving genes are more divergent than slowly evolving genes. Therefore, for these factors, patterns opposite to the reality are expected: the fastest-evolving dataset has the largest *SumT*/*SumI* value, and the slowest-evolving dataset has the smallest *SumT*/*SumI* value. Besides, mitigating the rate heterogeneity or selection heterogeneity among sites can actually aggravate the extension of the terminal branches. Take bin #30 as an example. The 3^rd^ positions of bin #30 has a rate approximately 10 times that of the 2^rd^ positions. Some rate heterogeneity or selection heterogeneity is apparent in bin #30. As mentioned above, concatenating all sites of bin #30 as one partition did not show a prominent extension of the terminal branches. In comparison, using only the 2^nd^ position would make the dataset less heterogeneous, which did not alleviate the extension of the terminal branches but, instead, aggravated it. Thereby, selection heterogeneity among sites and rate heterogeneity among sites are also unlikely to explain the extension of the terminal branches.

Additionally, we investigated whether some other factors can explain the extension of the terminal branches (see Supplementary Methods). First, we analyzed whether the relative composition variability (RCV) can explain the extension of the terminal branches (Phillips and Penny 2003). We investigated the correlation between RCV and *ω*. We found that the RCV value is negatively correlated with *ω* (Figure S6A, left). However, when we regrouped the 2242 coding sequences into 30 bins by RCV values, the branch length patterns for the three codon positions (Figure S6A, right) were different from those in Figure 4. Thus, RCV is unlikely to be responsible for the extension of the terminal branches. Additionally, we analyzed whether the GC content can explain the extension of the terminal branches. We investigated the correlation between the mean GC content of gene and *ω*. We found that the mean GC content is positively correlated to *ω* (Figure S6B, left). When we regrouped the 2242 CDS into 30 bins by GC content, although we observed a pattern slightly homologous to the extension of the terminal branches (Figure S6B, right), that pattern is far less prominent than the pattern that we have shown above (Figure 4). Thus, the GC content is also unlikely to be responsible for the extension of the terminal branches. Gene tree discordance can also influence the inference of branch lengths (Mendes and Hahn 2016). However, gene tree discordance is expected to influence the length of the whole tree rather than just terminal branches or internal branches. Furthermore, this impact is generally modest. Thus, gene tree discordance seems also to be implausible for explaining the extension of the terminal branches. For now, the TDMR caused by purifying selection seems to be a more reasonable explanation for the extension of the terminal branches rather than other factors.

In an influential study about TDMR, Ho et al. (2005), the authors depicted trends of rates against time for three cases: mitochondrial protein-coding genes of avian taxa, mitochondrial protein-coding genes of primates and D-loop sequences of primates. In Ho et al. (2005), the authors claimed that the TDMR trends reached plateaus before 2 Ma. According to Ho et al. (2005), the TDMR caused by purifying selection seems not able to influence the deep time scales involved in the present study. However, due to the limited data size, large uncertainties remain in the result of Ho et al. (2005), the point of reaching the plateau can be also 5, 6, or even 10 Ma (Woodhams 2005). More importantly, the result of Ho et al. (2005) was based on all sites of genes. In the present study, when concatenating all sites of genes as one partition, the extension of the terminal branches is actually not prominent. Nevertheless, the time depth that is influenced by the TDMR caused by purifying selection depends on the selective constraint. In a previous study, Subramanian and Lambert (2011), the authors compared the TDMR trends of the nonsynonymous data and the synonymous data for mitochondrial genes of humans and chimpanzees. For the synonymous data, before 10 Ma, the trend had reached the plateau, whereas for nonsynonymous data, until 10 Ma, the trend had not yet reached the plateau. This result suggests that the stronger the selective constraint is, the greater time depth is influenced by the TDMR caused by purifying selection. Hence, simply from studies based on sites under the average selective constraint, we should not conclude that the TDMR caused by purifying selection cannot influence species-level molecular dating. Moreover, the result of Ho et al. (2005) was based on mitochondrial genes. Mitochondrial genomes have smaller effective population sizes than nuclear genomes. The fixation time for mitochondrial genes is expected to be shorter than nuclear genes. Thus, purifying selection could influence a deeper timescale for nuclear genes than for mitochondrial genes. Attributing the extension of the terminal branches to the TDMR caused by purifying selection is not conflict with the existing empirical evidences.

However, the theoretical studies based on the Wright-Fisher model suggest that large effective population sizes are required to explain the TDMR pattern observed in Ho et al. (2005) by purifying selection alone (Woodhams 2005; O’Fallon 2010). There exist a disparity between the theoretical evidences and the empirical evidences. Thus, finding a perfect explanation for the extension of the terminal branches seems to be a puzzle. In spite of this, as discussed above, the TDMR caused by purifying selection shows a different explanatory ability for the extension of the terminal branches, using other factors to explain why the extension of terminal branches depends on the selective constraint is difficult. Hence, on present evidence, the TDMR caused by purifying selection seems, at least, to be an important contributor to the extension of the terminal branches.

### The Implication for Molecular Dating Practices

In this study, we observed that, as the selective constraint becomes stronger, terminal branches are relatively extended. Although it is difficult to find a perfect explanation for this result, the result itself implies that purifying selection has an impact on species-level molecular dating. In population-level molecular dating, some researchers have suggested using selectively neutral genes/sites to avoid the impact of purifying selection (Subramanian et al. 2009; Subramanian and Lambert 2011, 2012). Similarly, for the species-level case in this study, such a method should also be recommended.

On the other hand, as mentioned in the Introduction, in current practices of species-level molecular dating, researchers would like to select slow-evolving genes/sites to reduce the impact of substitution saturation. These researchers may believe that the only disadvantage of excluding fast-evolving genes/sites is the reduction of the information content; no bias would be introduced by this method. From this perspective, if the dataset is large enough, the selection of slow-evolving genes/sites seems to be more elaborate and reliable (dos Reis et al. 2012; Jarvis et al. 2014). In the present study, from the result of the 1P scheme in Figure 6 and the comparison among the 3^rd^ position and all sites in the randomly sampled genes (Figure 7), we can see that if we do not intentionally select some genes/sites, purifying selection would not dramatically influence the time estimate in the species-level molecular dating. However, the selection of slow-evolving genes/sites can strengthen the impact of purifying selection. In extremes, the impact of purifying selection can be strengthened so much that it biases the time estimate dramatically (e.g., the result based on the 2^nd^ position of the slowest genes). If one prefers to select slowly evolving gene/sites, the result could be misleading. Thus, the opinion that selecting slow-evolving genes/sites cause no harm to the accuracy of species-level molecular dating may need to be reconsidered.

Nevertheless, our study does not mean that there is no need to avoid substitution saturation. It is reasonable to remove those genes/sites with exceptionally fast rates from data because the fast rates of these genes/sites may result from positive selection or mutational hotspots (Pisani 2004; Zheng et al. 2004). Additionally, in some cases, such as using mitochondrial genes or/and estimating highly deep divergences, selecting genes/sites under relaxed selective constraints may increase the risk of being influenced by substitution saturation, and using those genes/sites with slower rates may be more reasonable. Hence, through considering the impact of purifying selection, a question is raised: How can a trade-off be made between avoiding purifying selection and avoiding substitution saturation? Further studies are required to address this question. With further studying of this question in the future, researchers may be able to get more reliable results in species-level molecular dating. All in all, in species-level molecular dating, the impact of purifying selection should not be neglected.

