## Supplementary Table 1 for "Evaluating the Impact of Purifying Selection on Species-level Molecular Dating"

**Table S1. The result of testing whether the number of linear regression trends that show positive slope values has a statistical significance.** We define the “success” as that the linear regression trend has a positive slope value, and the “failure” the linear regression trend has a non-positive slope value. In each cell, the number of “successes” and the number of “failures” are shown for each comparison between two treatments. When *p*<0.05, the number of “successes” is significantly greater than the number of “failures”.

|  | Number of success | Number of failures | *p*-value |
| --- | --- | --- | --- |
| Branch length, 1^st^ positions | 23 | 0 | 1.91×10^-7^ |
| Branch length, 2^nd^ positions | 23 | 0 | 1.91×10^-7^ |
| Branch length, 3^rd^ positions | 15 | 8 | 0.05 |
| Time, 1^st^ positions | 20 | 0 | 9.53×10^-7^ |
| Time, 2^nd^ positions | 18 | 2 | 2.01×10^-4^ |
| Time, 3^rd^ positions | 18 | 2 | 2.01×10^-4^ |
| Time, All positions, 1P | 19 | 1 | 2.0×10^-5^ |
| Time, All positions, 3P | 20 | 0 | 9.53×10^-7^ |
