## Supplementary Table 2 for "Evaluating the Impact of Purifying Selection on Species-level Molecular Dating"

| 3^rd^ | All sites | 1^st^ | 1^st^ + 2^nd^ | 2^nd^ |  |
| --- | --- | --- | --- | --- | --- |
| - | *t* = -21.2,  *p* = 7.95×10^-53^ | *t* = -31.25,  *p* = 1.73×10^-78^ | *t* = -36.79,  *p* = 1.88×10^-90^ | *t* = -40.45,  *p* = 1.15^-97^ | 3^rd^ |
|  | - | *t* = -15.18,  *p* = 5.06×10^-35^ | *t* = -21.97,  *p* = 5.51×10^-55^ | *t* = -28.11,  *p* = 4.81×10^-71^ | All sites |
|  |  | - | *t* = -6.93,  *p* = 5.76×10^-11^ | *t* = -14.65,  *p* = 2.06×10^-33^ | 1^st^ |
|  |  |  | - | *t* = -8.21,  *p* = 2.86×10^-14^ | 1st + 2nd |
|  |  |  |  | - | 2nd |

**Table S2. The result of testing whether the extension of the terminal branches observed in the random sampling comparison has a statistical significance.** In each cell, the mean difference in *SumT*/*SumI* between two treatments (*t*-value) is shown. A negative value means the dataset under a more relaxed constraint has a smaller *SumT*/*SumI* value. When *p*<0.05, the result is statistically significant.
