## Supplementary Table 3 for "Evaluating the Impact of Purifying Selection on Species-level Molecular Dating"

| 3^rd^ | All sites | 1^st^ | 1^st^ + 2^nd^ | 2^nd^ |  |
| --- | --- | --- | --- | --- | --- |
| - | 2065:135,  *p* = 0.0 | 2051:149,  *p* = 0.0 | 2055:145,  *p* = 0.0 | 2066:134,  *p* = 0.0 | 3^rd^ |
|  | - | 1822:378,  *p* = 3.21×10^-226^ | 1994:206,  *p* = 0.0 | 1999:201,  *p* = 0.0 | All sites |
|  |  | - | 1823:377,  *p* = 6.65×10^-227^ | 1756:444,  *p* = 4.93×10^-184^ | 1^st^ |
|  |  |  | - | 1597:603,  *p* = 2.19×10^-184^ | 1st + 2nd |
|  |  |  |  | - | 2nd |

**Table S3. The result of testing the impact on time estimates for the random sampling comparison.** We define the “success” as the dataset under a strong constraint gives an older time estimate, and the “failure” as the dataset under a strong constraint does not give an older time estimate. In each cell, the number of “successes” and the number of “failures” are shown for each comparison between two treatments. When *p*<0.05, the number of “successes” is significantly greater than the number of “failures”.
