## Supplementary Table 4 for "Evaluating the Impact of Purifying Selection on Species-level Molecular Dating"

| Table S4. Substitution saturation test based on Xia's test using DAMBE | | | | | | | | | |
| --- | --- | --- | --- | --- | --- | --- | --- | --- | --- |
|  | 1st | | | 2nd | | | 3rd | | |
| subset | Iss | Iss.c | *p* | Iss | Iss.c | *p* | Iss | Iss.c | *p* |
| 1 | 0.4006 | 0.8497 | 0.0000 | 0.3655 | 0.8497 | 0.0000 | 0.4976 | 0.8497 | 0.0000 |
| 2 | 0.3524 | 0.8438 | 0.0000 | 0.3214 | 0.8438 | 0.0000 | 0.4780 | 0.8438 | 0.0000 |
| 3 | 0.3405 | 0.8459 | 0.0000 | 0.3052 | 0.8459 | 0.0000 | 0.4878 | 0.8459 | 0.0000 |
| 4 | 0.3194 | 0.8430 | 0.0000 | 0.2840 | 0.8430 | 0.0000 | 0.4743 | 0.8430 | 0.0000 |
| 5 | 0.3012 | 0.8399 | 0.0000 | 0.2631 | 0.8399 | 0.0000 | 0.4627 | 0.8399 | 0.0000 |
| 6 | 0.3153 | 0.8392 | 0.0000 | 0.2791 | 0.8392 | 0.0000 | 0.4815 | 0.8392 | 0.0000 |
| 7 | 0.2780 | 0.8347 | 0.0000 | 0.2446 | 0.8347 | 0.0000 | 0.4507 | 0.8347 | 0.0000 |
| 8 | 0.2958 | 0.8356 | 0.0000 | 0.2589 | 0.8356 | 0.0000 | 0.4797 | 0.8356 | 0.0000 |
| 9 | 0.2961 | 0.8348 | 0.0000 | 0.2618 | 0.8348 | 0.0000 | 0.4714 | 0.8348 | 0.0000 |
| 10 | 0.2604 | 0.8291 | 0.0000 | 0.2283 | 0.8291 | 0.0000 | 0.4439 | 0.8291 | 0.0000 |
| 11 | 0.2466 | 0.8288 | 0.0000 | 0.2140 | 0.8288 | 0.0000 | 0.4318 | 0.8288 | 0.0000 |
| 12 | 0.2446 | 0.8279 | 0.0000 | 0.2109 | 0.8279 | 0.0000 | 0.4456 | 0.8279 | 0.0000 |
| 13 | 0.2445 | 0.8298 | 0.0000 | 0.2116 | 0.8298 | 0.0000 | 0.4389 | 0.8298 | 0.0000 |
| 14 | 0.2345 | 0.8234 | 0.0000 | 0.2009 | 0.8234 | 0.0000 | 0.4309 | 0.8234 | 0.0000 |
| 15 | 0.2329 | 0.8264 | 0.0000 | 0.2023 | 0.8264 | 0.0000 | 0.4296 | 0.8264 | 0.0000 |
| 16 | 0.2265 | 0.8243 | 0.0000 | 0.1971 | 0.8243 | 0.0000 | 0.4287 | 0.8243 | 0.0000 |
| 17 | 0.2355 | 0.8236 | 0.0000 | 0.2040 | 0.8236 | 0.0000 | 0.4379 | 0.8236 | 0.0000 |
| 18 | 0.2306 | 0.8252 | 0.0000 | 0.1991 | 0.8252 | 0.0000 | 0.4416 | 0.8252 | 0.0000 |
| 19 | 0.2114 | 0.8223 | 0.0000 | 0.1806 | 0.8223 | 0.0000 | 0.4292 | 0.8223 | 0.0000 |
| 20 | 0.2224 | 0.8224 | 0.0000 | 0.1932 | 0.8224 | 0.0000 | 0.4334 | 0.8224 | 0.0000 |
| 21 | 0.2329 | 0.8250 | 0.0000 | 0.2039 | 0.8250 | 0.0000 | 0.4379 | 0.8250 | 0.0000 |
| 22 | 0.2090 | 0.8219 | 0.0000 | 0.1796 | 0.8219 | 0.0000 | 0.4325 | 0.8219 | 0.0000 |
| 23 | 0.1912 | 0.8219 | 0.0000 | 0.1651 | 0.8219 | 0.0000 | 0.4073 | 0.8219 | 0.0000 |
| 24 | 0.1969 | 0.8222 | 0.0000 | 0.1722 | 0.8222 | 0.0000 | 0.4102 | 0.8222 | 0.0000 |
| 25 | 0.1911 | 0.8219 | 0.0000 | 0.1653 | 0.8219 | 0.0000 | 0.4199 | 0.8219 | 0.0000 |
| 26 | 0.1869 | 0.8219 | 0.0000 | 0.1630 | 0.8219 | 0.0000 | 0.4014 | 0.8219 | 0.0000 |
| 27 | 0.1838 | 0.8219 | 0.0000 | 0.1593 | 0.8219 | 0.0000 | 0.4079 | 0.8219 | 0.0000 |
| 28 | 0.1813 | 0.8221 | 0.0000 | 0.1593 | 0.8221 | 0.0000 | 0.4028 | 0.8221 | 0.0000 |
| 29 | 0.1740 | 0.8219 | 0.0000 | 0.1528 | 0.8219 | 0.0000 | 0.3980 | 0.8219 | 0.0000 |
| 30 | 0.1658 | 0.8221 | 0.0000 | 0.1465 | 0.8221 | 0.0000 | 0.3788 | 0.8221 | 0.0000 |

| Interpretation of results: | | |
| --- | --- | --- |
|  | Significant Difference | |
|  | Yes (*p* < 0.05) | No (*p* ≥ 0.05) |
| Iss < Iss.c | Little saturation | Substantial saturation |
| Iss ≥ Iss.c | Useless sequences | Very poor for phylogenetics |
