## Supplementary Figures for "Evaluating the Impact of Purifying Selection on Species-level Molecular Dating"

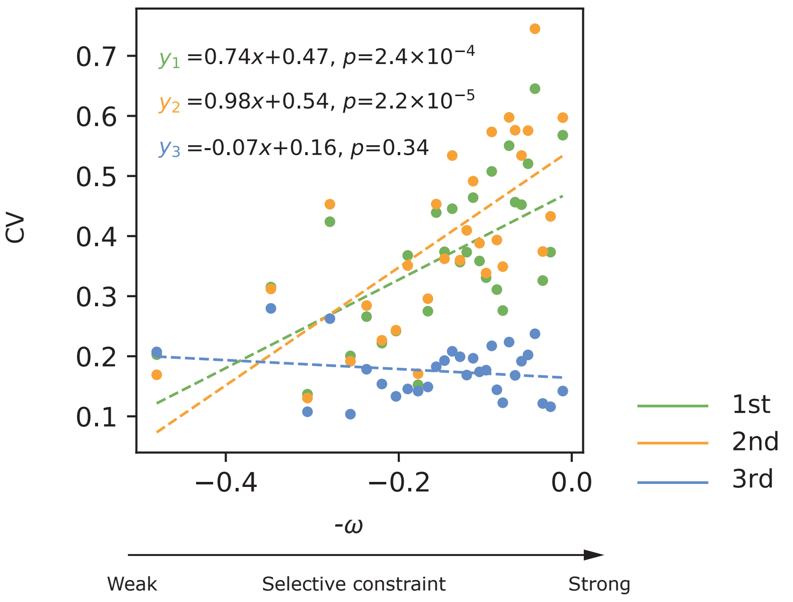


**Figure S1. The correlations between purifying seleciton and rate variation among Catarrhini ineages.** The *x*-axis is the opposite of the mean pairwise d*N*/d*S* (-ω), indicating the overall selective constraint on a bin (the right is under the stronger constraint). The *y*-axis is the coefficient of variation (CV) of node-to-tip distances across the Catarrhini, indicating the rate variation among Catarrhini lineages. Generally, as the selective constraint, the rate variation increases.

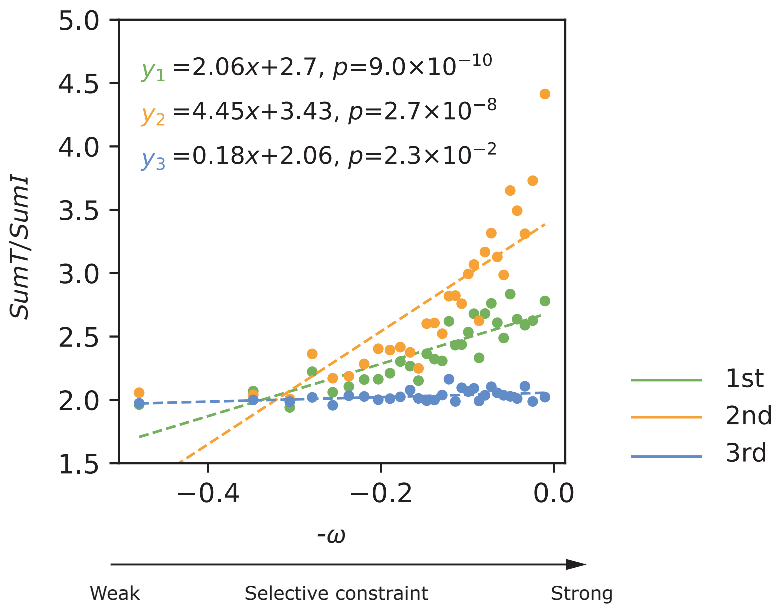


**Figure S2. The pattern of the branch lengths inferred by RAxML .** The *x*-axis is the opposite of the mean pairwise d*N*/d*S* (-ω), indicating the overall selective constraint on a bin (the right is under the stronger constraint). The *y*-axis is the ratio of the sum of terminal branch lengths to the sum of internal branch lengths (*SumT*/*SumI*), indicating the overall relative length of terminal branches. This result is similar to the result inferred by MCMCTree (see Fig. 4, upper).

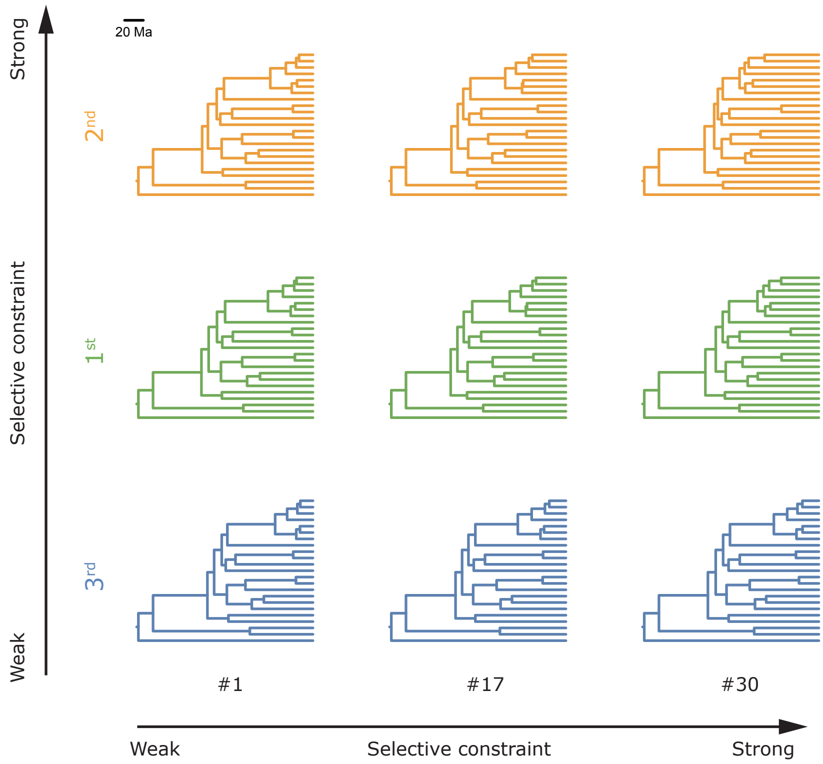


**Figure S3. The time trees of three representative bins.** Each time tree corresponds to a phylogram tree in Fig. 4. The time trees are inferred from different codon positions of three representative bins "#1", "#17" and "#30", which are under the least, moderate and strongest selective constraints, respectively. The topology of each tree follows that of Fig. 2.

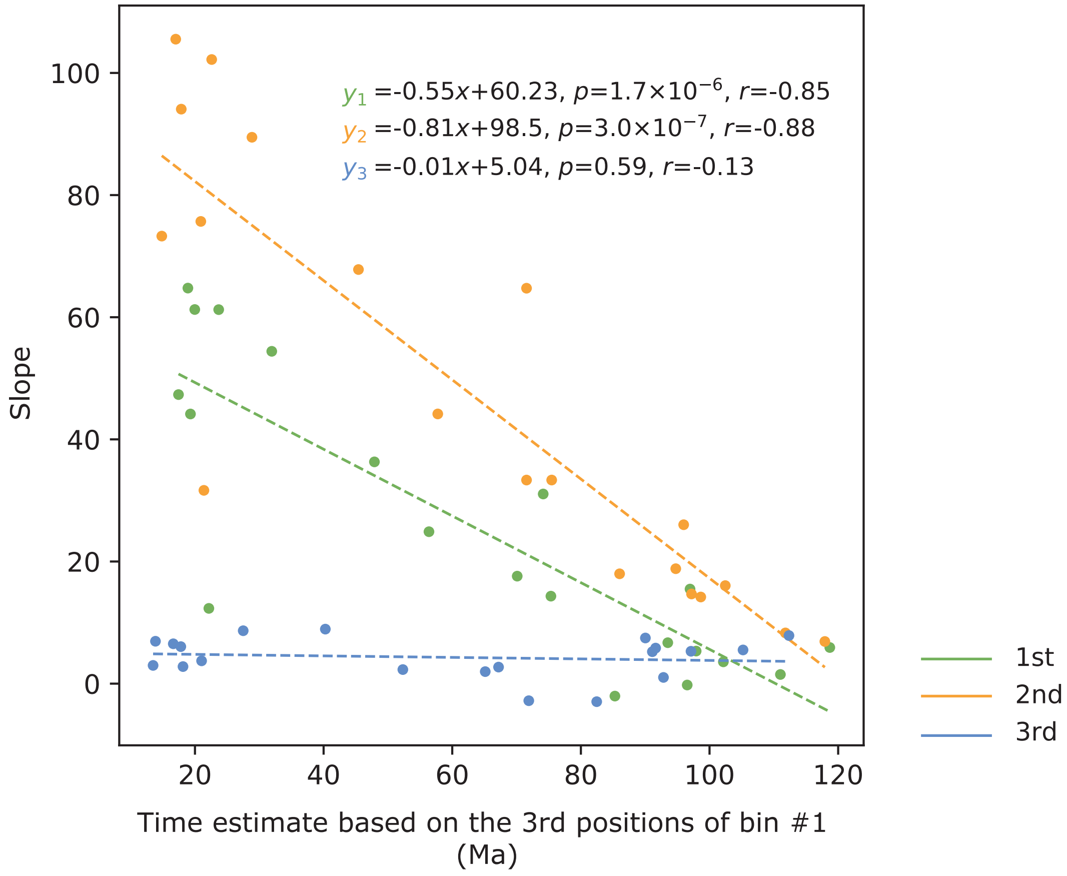


**Figure S4. The correlation between the slope and time depth.** The *x*-axis is the time estimate based on the 3rd positions of bin #1 (under the most relaxed constraint). The *y*-axis is the slope value shown in Fig. 6. The younger the divergence time is, the larger the slope value is.

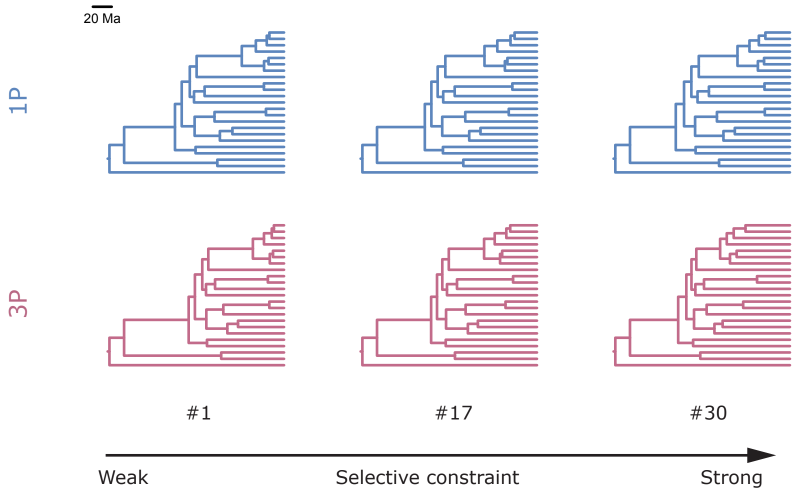


**Figure S5. The representative time trees for showing the impact of purifying selection under different partitioning schemes.** The time trees are inferred from three representative bins “#1”, “#17” and “#30”, which are under the least, moderate and strongest selective constraints, respectively. All the codon positions were used to infer the time trees here, two different partitioning schemes were adopted: concatenating all sites as one partition (1P) and partioning the dataset by codon positions (3P). Each node in each time tree corresponds to a dot in Fig. 7. The topology of each tree follows that of Fig. 2.

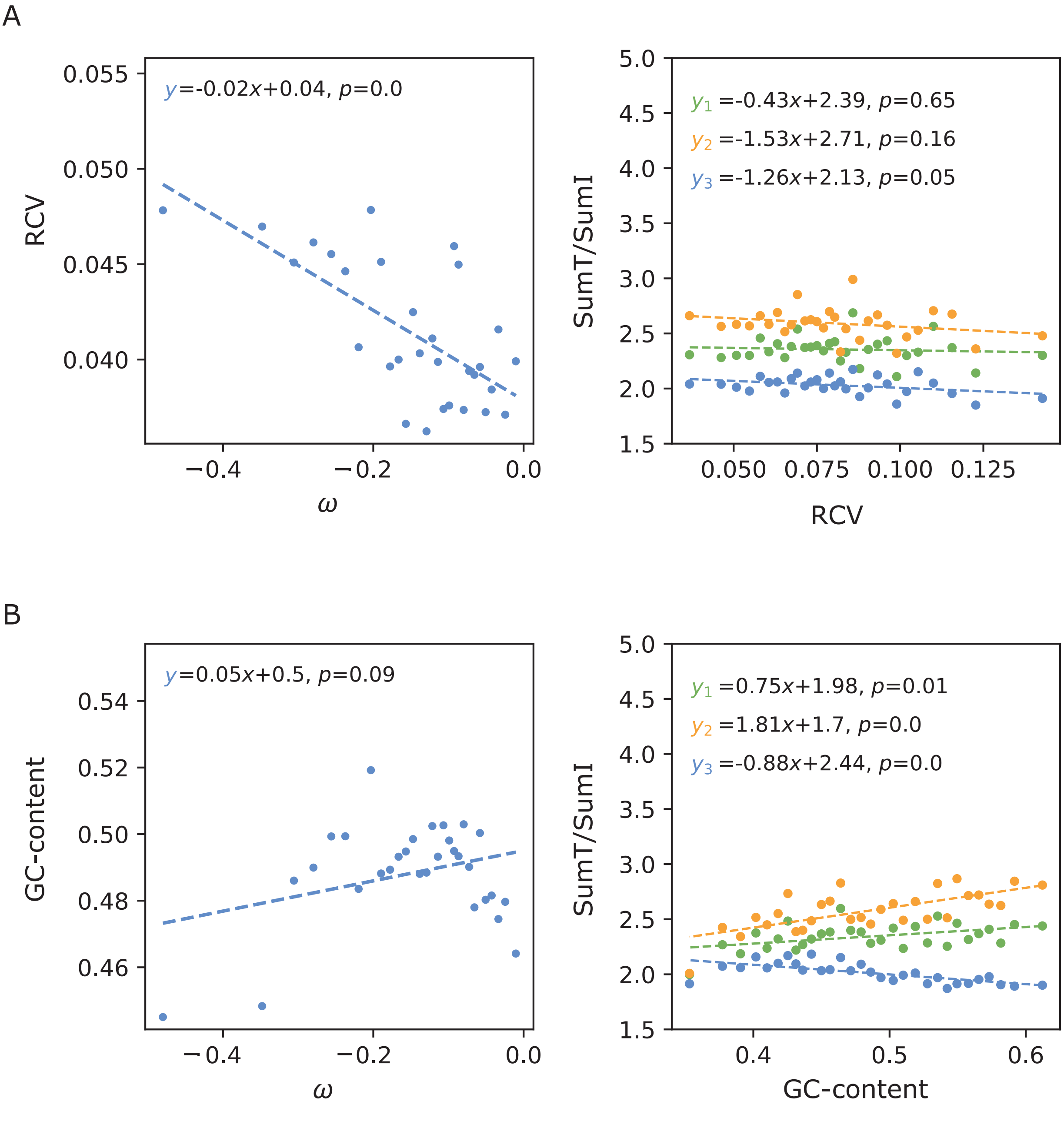


**Figure S6. A)** The correlation between RCV and the seletive constraint (left) and the pattern of the branch length for the bins grouped by RCV (right). **B)** The correlation between the GC-content and the seletive constraint (left) and the pattern of the branch length for the bins grouped by GC-content (right).
